# Interaction mechanism between the HSV-1 glycoprotein B and the antimicrobial peptide Amyloid-β

**DOI:** 10.1101/2022.02.17.480815

**Authors:** Karine Bourgade, Eric H. Frost, Gilles Dupuis, Jacek M. Witkowski, Benoit Laurent, Charles Calmettes, Charles Ramassamy, Mathieu Desroches, Serafim Rodrigues, Tamás Fülöp

## Abstract

Unravelling the mystery of Alzheimer’s Disease (AD) requires urgent resolution given the worldwide increase of the aging population. There is a growing concern that the current leading AD hypothesis, the amyloid cascade hypothesis, does not stand up to validation with respect to emerging new data. Indeed, several paradoxes are being discussed in the literature, for instance, both the deposition of the Amyloid-Beta peptide (Aβ) and the intracellular neurofibrillary tangles (NFTs) could occur within the brain without any cognitive pathology. Thus, these paradoxes suggest that something more fundamental is at play in the onset of the disease and other key and related pathomechanisms have to be investigated. The present study follows our previous investigations on the infectious hypothesis, which posits that some pathogens are linked to late onset AD. Our studies also build upon the shattering finding that Aβ is a powerful antimicrobial agent capable of inhibiting pathogens as observed in *in vitro* experiments. Herein, we ask what are the molecular mechanisms in play when Aβ neutralizes infectious pathogens? To answer this question, we probed at nanoscale lengths with FRET (Förster Resonance Energy Transfer), the interaction between Aβ peptides and glycoprotein B (responsible of virus-cell binding) within the HSV-1 virion. We concluded that there is indeed a close interaction, likely nonspecific or semi-specific, between the two types of molecules, which participate in virus neutralization.

## Introduction

Alzheimer’s disease (AD) is the leading cause of dementia in the world, mainly due to the global increase of the aging population, as age is one of the most important risk factors for developing AD. Despite intense research efforts, we still do not know what the exact causes of late onset AD are. Nevertheless, there is a general agreement that AD’s prime histopathological signature led to the so-called *Amyloid Cascade Hypothesis* (ACH), involving plaques induced by Amyloid-beta (Aβ) peptides and of intracellular neurofibrillary tangles (NFTs) deposition (Sanabria-Castro et al., 2017). However, several paradoxes forced us to expand this view of ACH and reconcile it with novel emerging data, e.g.: 1) Aβ peptide deposition can occur in cognitively normal individuals (Poul et al., 2020); 2) Aβ peptides are abundant in the mild cognitive impairment stage of AD and decrease in some clinically diagnosed AD (Fülöp et al., 2018); 3) inflammation precedes Aβ peptide deposition (Kinney et al., 2018); 4) current data support the view that aberrant processing of APP (amyloid precursor protein) towards Aβ peptides, may sometimes cause human familial/early onset AD. All current data do not support the conclusion that aberrant Aβ peptide expression is the cause of late onset AD but likely only plays a secondary role as part of a more complex process. A compelling (yet controversial) body of data is mounting, which can be reconciled with ACH and potentially explain the above paradoxes. This data has its origin in a hypothesis by Dr O. Fischer in 1910, which posited that infectious pathogens are involved in AD since senile plaques are reminiscent of bacterial colonies (Fischer, 1910). Independent research (including ours) has been instrumental in corroborating the infectious hypothesis by showing that, in fact, a family of microorganisms (e.g. HSV-1, spirochetes, *P. gingivalis*) are potentially associated with late onset AD, also hinting that sporadic AD is a syndrome (Fülöp et al., 2018; Itzhaki, 2004). As a case in point, DNA of HSV-1 has been found in senile plaques (Lövheim et al., 2015). Moreover, the demonstration by several groups that the Aβ peptide is a powerful antimicrobial peptide secreted by neurons in response to attacks by microorganisms. This demonstration called Amyloid Protection Hypothesis (APH) lends weight to the infection hypothesis (IH) (Bourgade et al., 2016; Vijaya et al., 2016). Thus, under APH-IH theory, Aβ peptide accumulation is no longer seen as Aβ being the main participant in the pathophysiology of AD, but rather as an innate immune response (i.e. reversing its role). Thus, senile plaques are possibly a by-product of Aβ fighting to contain and neutralize infections.

As HSV-1 is a recognized culprit to foment APH/IH, its interaction with Aβ is of considerable interest. Initially it was demonstrated that APP, the precursor of Aβ, associated intracellularly with HSV-1 and contributed to the movement of the virus towards the surface (anterograde transport) (Satpute-Krishnan et al., 2006, 2011). It has been shown that pathogens and particularly HSV-1, interfere with the APP metabolic pathway and use C99 protein (specifically, the final 15 amino-acid sequence), rather than Aβ, to coopt the intracellular transport machinery, enabling their transport along microtubules, but also causing NFTs (Satpute-Krishnan et al., 2006, 2011; Cheng et al., 2011). Moreover, studies demonstrate that APP C-terminal fragment, C99 protein, but not Aβ peptides is associated with neuronal death (Pulina et al., 2019).

The interaction between viruses and amyloid peptides is under intense investigations to confirm the pathogenic role of viruses in various diseases, including neurodegenerative diseases. In most cases, the interaction is studied as the putative cause of the viral pathogenesis by inducing amyloid aggregations. Recently, the interaction between viral proteins including HSV-1 and the Aβ42 peptide has been described in different setting and aiming to take advantage that nanoparticles have been shown to act as catalytic surfaces that facilitate heterogenous nucleation of amyloid fibrils *via* binding, concentrating, and enabling conformational changes of amyloidogenic peptides (Idrees and Kumar, 2021; Hsu et al., 2021; Hensel et al., 2020). In 2019, Ezzat *et al*., have demonstrated that HSV-1 accelerated the kinetics of Aβ42 peptides aggregation and to a lesser extent that of Aβ40 peptides. The authors have demonstrated an interaction between amyloid fibrils and the viral surface at different stages of maturation *via* early protofibrillar intermediates. They also observed that HSV-1 infection led to increased accumulation of amyloid plaques in a mouse model of AD. In their discussion, the authors try to reconcile the antimicrobial characteristics and the virus induced nucleation process, leading to amyloid plaques. Certainly, this is not mutually exclusive as we have shown that, at the beginning, the Aβ42 peptide secreted by cells is antimicrobial but as the infection is progressing, it becomes pathogenic, by the virtue of the process described by Ezzat *et al*. This reminds us of the capacity of bacteria to form biofilm which mimic plaques and the fact that biofilms have already been demonstrated in AD (Miklossy, 2016). These studies focus only on the putative pathogenic interaction of HSV-1 and Aβ42 peptides, resulting in amyloid plaque formation, rather than the antimicrobial interaction of Aβ42 against HSV-1. Indeed, these studies have presented corona formation as an essential step in the viral infection.

We have demonstrated earlier that the Aβ peptide, as antimicrobial peptide, prevents HSV-1 infection (Bourgade et al., 2015, 2016). Indeed, we have demonstrated that Aβ peptides addition in the culture media before or at the same time as HSV-1, decreases virus infectivity, while adding Aβ peptides when HSV-1 has already entered the neurons, show no effects (Bourgade et al., 2015). Moreover, HSV-1-infected neurons supernatants, when added to new neuronal cultures, were also able to inhibit infection by HSV-1, presumably due to induced production of Aβ (Bourgade et al., 2016). This accumulating evidence has led us to ask about the molecular mechanisms of actions of Aβ peptides against AD pathogens.

Taking these data into consideration, it appears highly important to explore the currently unknown mechanism whereby Aβ peptides render HSV-1 non-infectious. This potential gB-Aβ peptides interaction could also be extended to other enveloped viruses, as well as peptides, and could represent a threshold when it becomes inefficient and pathological. Thus, the aim of the present study was to gather more experimental data to determine the antiviral mechanism of Aβ peptides against HSV-1 virus.

## Material and Methods

### Sequence alignment and antimicrobial prediction

The amino acid sequence alignment between HSV-1 gB and Aβ42 was performed using the PyMol software after having depicted the 3D structure of HSV-1-gB and Aβ42 peptide (The PyMOL Molecular Graphics System, Version 2.0 Schrödinger, LLC (Bramucci et al., 2012)).

AntiBP2 software was used, with neural networks and support vector machines (SVM) to predict the amino-acid sub-sequence for a peptide with antibacterial activity. AntiBP2 utilizes four datasets to train their models: N-terminus based, C-terminus based, N+C terminus based and amino acid composition methods. These 4 methods are SVM trained on 4 different datasets, compiled using N, C, NC and full composition peptides respectively. For antiviral activity predictions, we employed the AVPpred software, which computes various features (i.e. motifs and alignments, followed by amino acid composition and physicochemical properties), during 5-fold cross validation using SVM. In particular, we have fragmented the amino sequence into subsequences of lengths 15, while taking the overlap length to be 14 and finally the subsequences of length 15 were processed by AVPred (Thakur et al., 2012).

### Förster Resonance Energy Transfer (FRET)

The Förster Resonance Energy Transfer (FRET) has been performed in a LightCycler480 II (Roche, Canada) with variable excitation and detection filters. 1 μg of a recombinant HSV-1 containing gB fused to the green fluorescent protein (GFP) (HSV-1-gB-GFP) produced at the Cochin Institute, Paris, France and kindly provided by Pr. Flore Rozenberg was mixed with 1 μg of Aβ42-HiLyteFluor-555 (Anaspec, Fremont, CA), with or without various pre-treatments (see below) and controls, without cells, directly in wells of a 96 well plate at room temperature. Plates were sealed, centrifuged, and then processed by the thermocycler. Thermocycler conditions were heating 5 min to reach 37°C, then fluorescence acquisition every 10 min for 30 min, before cooling. Fluorescence acquisition was measured with excitation at 440 nm and emission at 580 nm (green fluorescence) and also with excitation at 440 nm and emission at 660 nm (red fluorescence). Several controls were also tested with the same program. Pre-treatments included mixing Aβ42-HiLyteFluor-555 with 1 μg of blocking antibodies (α-Aβ42-Ab, Anaspec, Fremont, CA) for 15 min at room temperature prior to adding viral particles or pretreating the HSV-1-gB-GFP for 30 min at room temperature with 1% NP40 to disrupt the viral envelope, before mixing with Aβ42-HiLyteFluor-555. Controls included negative controls without virus or Aβ42-HiLyteFluor-555, virus alone, and Aβ42-HiLyteFluor-555 alone.

At the end of the program, the thermocycler software gave fluorescence values for green and red fluorescence for each well. Data obtained for each measure were averaged and used to create the graph.

Statistical analysis was performed using two-tailed unpaired T-test.

## Results

We first depict the 3D structure of HSV-1-gB and Aβ42 peptides (Fig. 1A). Sequence homology was observed between the viral gB protein and Aβ42 (Fig. 1B and Bourgade et al., 2016). Furthermore, the antibacterial predictions show that a subsequence containing the helix and C-terminus of Aβ42 peptide has antiviral activity (Fig. 1C and Ezzat et al., 2019). This result prompted us to analyze the interaction between HSV-1 and Aβ peptides on *in vitro* experiments.

**Figure 1.**
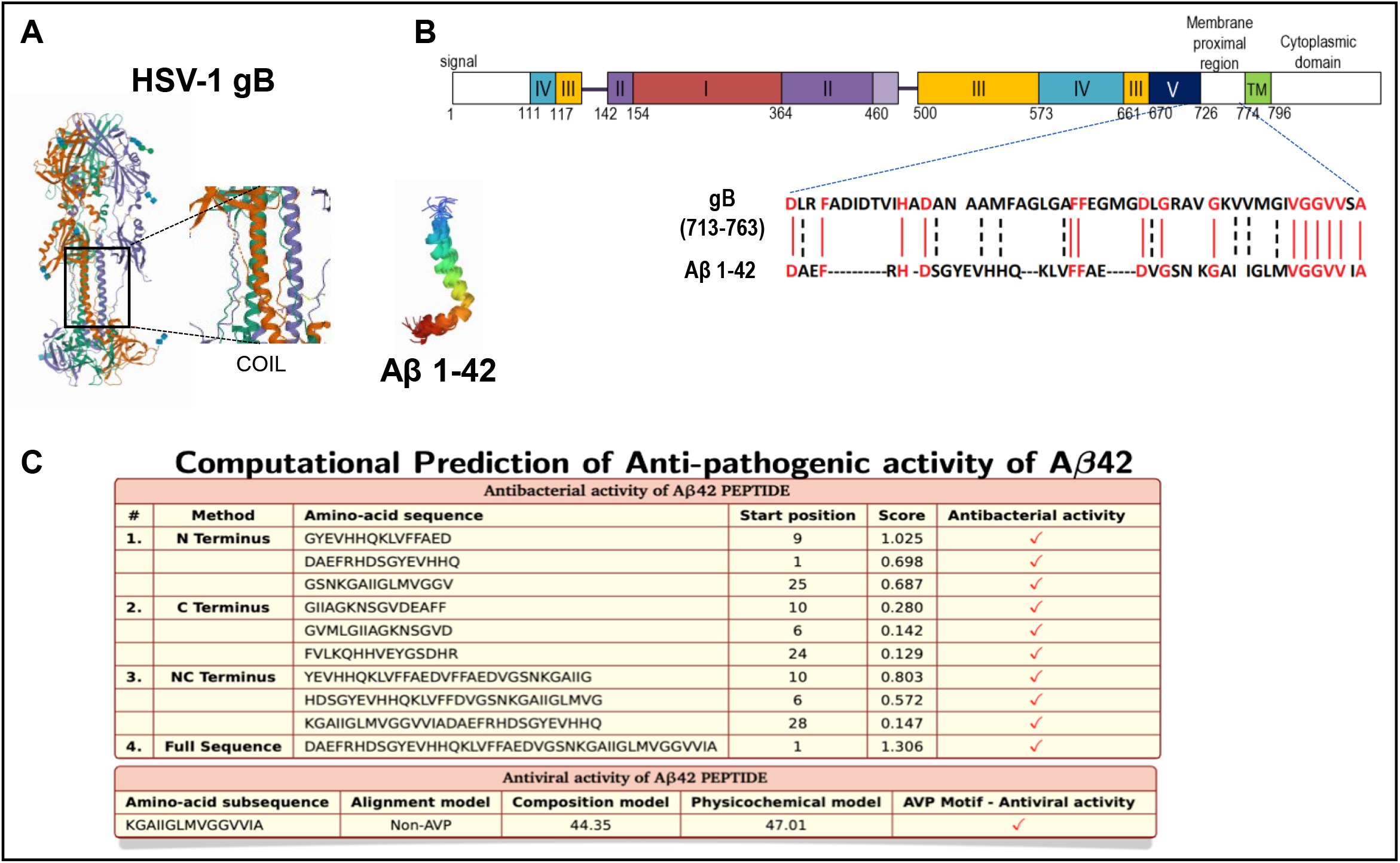
Molecular structure, sequence alignment and anti-pathogenic property of Aβ. Panel (A): 3D crystal structure of HSV-1-gB and Aβ-42; data from PDB (RCSB Protein Data Bank, http://www.rcsb.org) are shown using PyMol software. Center panel shows zoom on HSV-1 gB intermembrane sequence with, formed coils. Panel (B): sequence alignment between HSV-1-gB and Aβ-42 (using PyMol software). Panel (C): prediction of amino-acid sub sequences of Aβ42 possessing antibacterial activity *via* AntiBP2 software (http://crdd.osdd.net/raghava/antibp2) and antiviral activity with AVPpred software (http://crdd.osdd.net/servers/avppred).

FRET experiments were performed between an HSV-1 particle with gB linked to the green fluorescent protein (HSV-1-gB-GFP) and fluorescents peptides (Aβ42-HiLyteFluor-555 (red)). Their analysis has shown that its functionality remains intact despite the presence of the GFP (Potel et al., 2002). FRET experiments are a well-established way to determine whether two molecules with complementary green and red fluorescent labels are in close interaction (≤ 10 nm) (Broussard et al., 2013). When the green fluorescent label is excited its emission can be transferred and excite a nearby red fluorescent label. These experiments usually employ fluorescence microscopy, but they also could be performed with thermocyclers equipped with laser induction and wavelength specific detectors. They measure fluorescence amplification created by dyes and can detect FRET when different fluorescents dyes are brought close together (Zeng et al., 2009; Martinez-Serra et al., 2014). If only green fluorescence is observed (excitation at 440 nm and emission at 580 nm) then no energy transfer has occurred. This would be expected when virus is tested alone or if the gB-GFP was more than 10 nm from the red dye linked to Aβ42 peptides. If red fluorescence is measured in the channel with excitation at 440 nm (which does not efficiently excite red fluorophores) and emission at 660 nm, then GFP has transferred its energy to HiLyteFluor-555, which can then emit giving values that are high on red axis and low on the green axis because the green fluorescence has been quenched by transfer to the red fluorophore. The observed values are presented in Fig. 2 and then analyzed statistically in Table 1.

**Figure 2.**
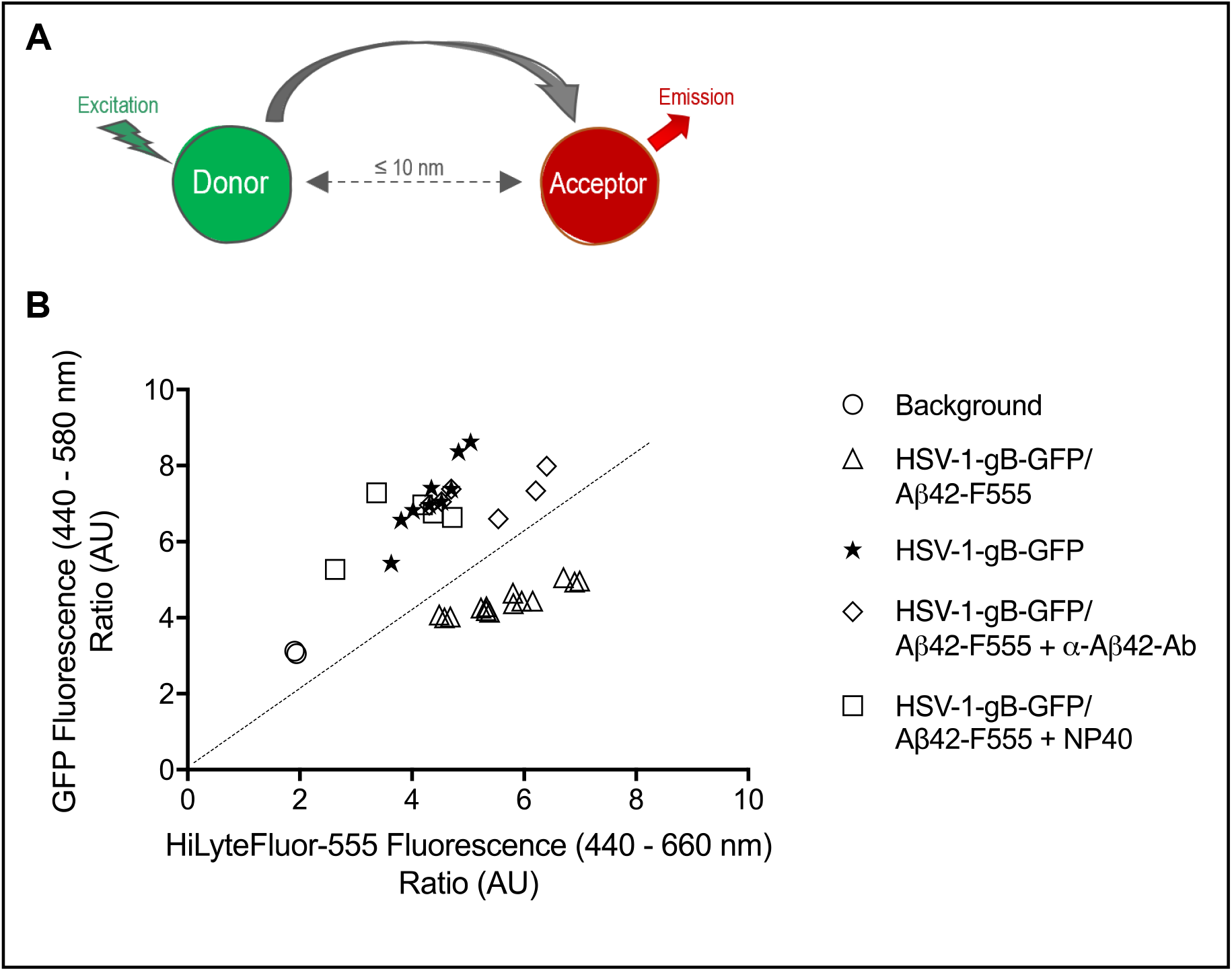
Energy transfer between HSV-1-gB-GFP and Aβ42-HiLyteFluor-555. Panel (A): Förster Resonance Energy Transfert process. Panel (B): ratio of emitted green and red fluorescence in each well. Green fluorescence was emitted by HSV-1-gB-GFP particles and red fluorescence was emitted by Aβ42-HiLyteFluor-555 when FRET occurs. Circles represent background measure, triangles represent HSV-1-gB-GFP/Aβ42-HiLyteFluor-555 mix, stars represent HSV-1-gB-GFP particles alone, squares show results after NP40 pre-treatment of virus and diamonds represent results after pre-treatment of Aβ42 with blocking antibodies. The dotted line identifies where green and red fluorescence emission are equal. n=2-5 independent experiments in triplicate.

**Table 1.**
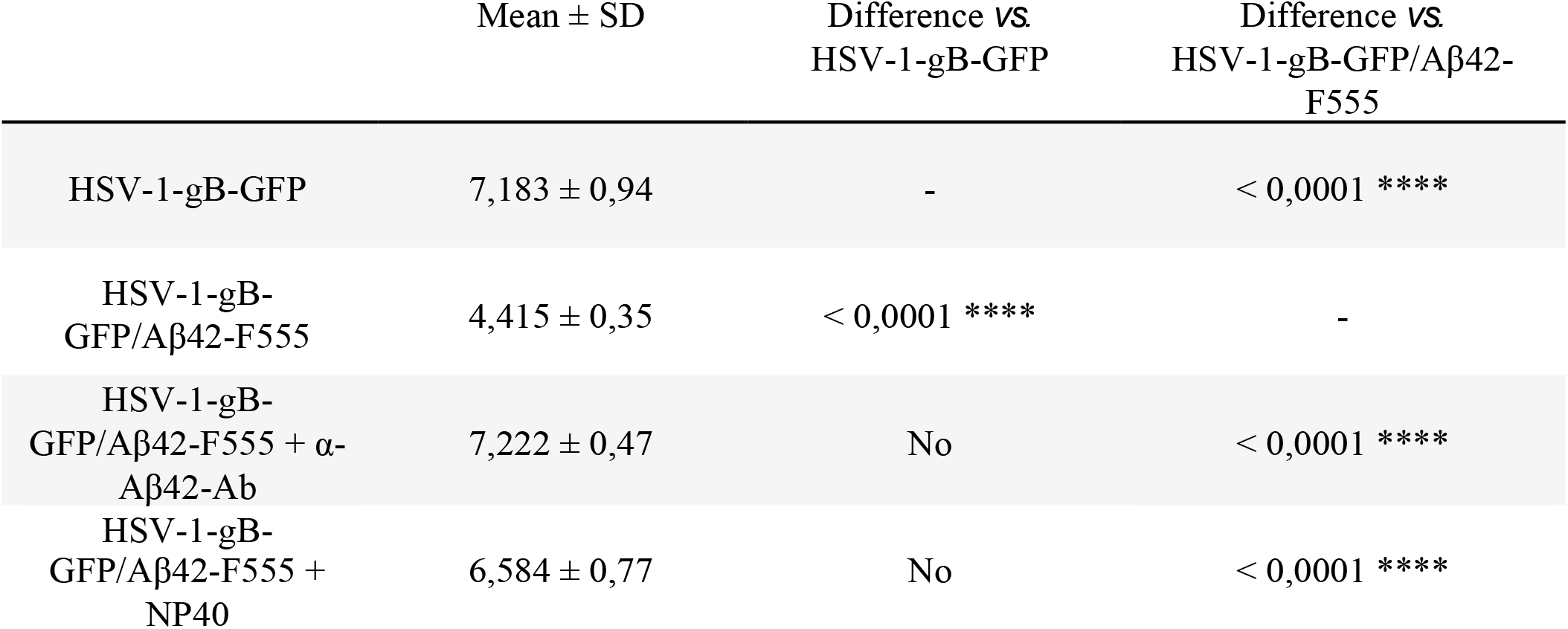
Documentation of FRET between HSV-1-gB-GFP and Aβ42-HiLyteFluor-555. Significant differences were observed in the mean fluorescence between HSV-1-gB-GFP/Aβ42-HiLyteFluor-555 mix and HSV-1-gB-GFP, but also when Aβ42-HiLyteFluor-555 was pre-treated with blocking antibodies or HSV-1-gB-GFP was pre-treated with NP40 prior to mixing. Unpaired T-test, two-tailed. ****: p<0,0001. n=2-5 independents experiments in triplicate.

| | Mean ± SD | Difference vs.<br>HSV-1-gB-GFP | Difference vs.<br>HSV-1-gB-GFP/Aβ42-F555 |
| --- | --- | --- | --- |
| HSV-1-gB-GFP | 7,183 ± 0,94 | - | < 0,0001 **** |
| HSV-1-gB-GFP/Aβ42-F555 | 4,415 ± 0,35 | < 0,0001 **** | - |
| HSV-1-gB-GFP/Aβ42-F555 + α-<br>Aβ42-Ab | 7,222 ± 0,47 | No | < 0,0001 **** |
| HSV-1-gB-GFP/Aβ42-F555 +<br>NP40 | 6,584 ± 0,77 | No | < 0,0001 **** |

Data showed that the green fluorescence emitted by HSV-1-gB-GFP alone (7,18 ± 0,94 AU) was significantly higher than that emitted by the HSV-1-gB-GFP/Aβ42-HiLyteFluor-555 mix (4,41 ± 0,35 AU) (Table 1), whereas the mix gave much higher fluorescence in the red channel than the virus alone did. Indeed, the ratio of red fluorescence divided by green fluorescence was superior to 1 for the HSV-1-gB-GFP/Aβ42-HiLyteFluor-555 mix, but inferior to 1 for HSV-1-gB-GFP alone. These results are consistent with FRET. We also observed that when HSV-1-gB-GFP was pre-treated with NP40 prior to mixing or when blocking antibodies directed against Aβ (α-Aβ42-Ab) were pre-mixed with Aβ42-HiLyteFluor-555, then mean green fluorescence values were 7,22 ± 0,47 AU and 6,58 ± 0,77 AU respectively, which are significantly different from the mean values without pre-treatment (4,41 ± 0,35 AU), and similar to those obtained for HSV-1gB-GFP alone. Furthermore, the ratio of red to green fluorescence also decreased to values lower than 1 in the pre-treated samples. These latter results corresponded to an absence of FRET.

## Discussion

The purpose of the FRET experiments carried out in this study was to determine the mechanism of action of Aβ peptides against HSV-1 virus. The present results indicated that the fluorescent energy resulting from the excitation of the GFP associated with the gB of HSV-1 virions was transmitted to the fluorochrome HiLyteFluor-555 associated with the peptide Aβ42. We concluded that FRET took place between the two fluorochromes, indicating that they are at a distance less than or equal to 10 nM (Broussard et al., 2013). Considering the proximity of Aβ42 and gB validated by FRET, the diameter of the HSV-1 virion which measures approximately 200 to 250 nm (Bohannon et al., 2013; Beilstein et al., 2019), and the diameter of HSV glycoprotein tetramers are probably close to 10 nm (glycoprotein D has been measured (Pilling et al., 1999)), so we can therefore conclude that the Aβ42 peptide must insert into the outer membrane of the virus, near the site on the viral membrane where gBs are present.

Controls show us that the energy transfer is prevented by the use of NP40 detergent, which cannot alter the peptides’ structure or direct interactions, suggesting that there is probably not a direct interaction between the peptide and the glycoprotein. The most logical explanation is that Aβ42 peptide is inserted into the viral envelope, near the gB. When the detergent has solubilized the viral envelope, insertion is prevented. Blocking antibodies also interfere with HSV-1-gB/Aβ42 interaction by preventing Aβ42 peptide from inserting itself into the viral envelope.

Our observations support the hypothesis that Aβ peptides insert into the outer membrane of enveloped viruses in the same way as the LL-37 peptide. LL-37 is an α helical peptide which inserts into lipid membranes and forms pores, deleterious to the integrity of the membrane and therefore to the organism or the target cell (Kai-Larsen and Agerberth, 2008; Turner et al., 1998). The Aβ42 peptide is also an α helix and studies on cells or bacterial membranes have shown that it is also able to insert into lipid bilayers and form pores (Lemkul and Bevan, 2013; Kagan et al., 2012). It is also possible that the Aβ42-HiLyteFluor-555 has integrated the corona as it has been shown to be present, but this would not explain the loss of infectivity. Indeed, it has been postulated that HSV-1 must be surrounded by a corona to be infectious.

Our results shown here seem to confirm (at least *in vitro*) that the Aβ acts as an intracerebral component of the humoral innate immune system. Thus, it would recognize some molecular patterns characteristic for the neurotropic viruses (including, but not limited to the HSV-1) in a non- or semi-specific way, possibly similar to the interactions between the PAMPs and Pathogen Recognition Receptors.

These experiments complement our previous and present results. When cells secret appropriate quantity of the Aβ42 peptides in the presence of enveloped viruses, this will initiate an interaction resulting in the neutralization of the virus infectivity. With age, the most important risk factor of AD, microglial clearance capacity decreases, and cellular and viral waste accumulate in brain. When the quantity of virus or the resulting synthesis of Aβ42 peptides increases, not only will the Aβ peptides inactivate the virus, but pathological interactions between the viruses e.g. HSV-1, RSV or SARS-CoV2 and Aβ peptides will result in the formation of traditional amyloid plaques. Upon further consideration, it can also be surmised that a biofilm or plaque may be advantageous both for the virus (protection) and for the cells (protection). The elucidation of these processes is of the most importance prior to using these conditions in a therapeutic perspective.

### Conclusion

We concluded that there is indeed a close interaction, likely nonspecific or semi-specific, between the two types of molecules, which result in virus neutralization. Aβ peptides are antimicrobial peptides (AMP) that belongs to innate immune response and insert into viral envelop to disrupt them and prevent infection.

## Acknowledgements

Special thanks to Pr. Flore Rozenberg at Institut Cochin (Paris, France) for kindly providing us the HSV-1-gB-GFP.

KB, EHF, GD and TF were supported by grants from the Canadian Institutes of Health Research (CIHR) (No. 106634), the Université de Sherbrooke, the Société des Médecins de l’Université de Sherbrooke (SMUS) and the Research Center on Aging.

SR was supported by Ikerbasque (The Basque Foundation for Science), by GV-AI-HEALTH, the Basque Government through the BERC 2018-2021 program, by the Spanish State Research Agency through BCAM Severo Ochoa excellence accreditation SEV-2017-0718 and through project RTI2018-093860B-C21 funded by (AEI/FEDER, UE) with acronym “MathNEURO”. MD and SR acknowledge the support of Inria *via* the Associated Team “NeuroTransSF”.

The authors declare no conflicts of interest.

